# ColFeatures: Automated data extraction and classification of bacterial colonies

**DOI:** 10.1101/2021.05.27.445853

**Authors:** Daniela A. García-Soriano, Frederikke Dybdahl Andersen, Jens Vinge Nygaard, Thomas Tørring

## Abstract

Examining microbial colonies on agar plates have been at the core of microbiology for many decades. It is usually done manually, and therefore subject to bias besides requiring a considerable amount of time and effort. In order to optimize and standardize the identification of bacterial colonies growing on agar plates, we have developed an open access tool available on GitHub: ColFeatures. The software allows automated identification of bacterial colonies, extracts key morphological data and generate labels that ensure tracking of temporal development. We included machine learning algorithms that provide sorting of environmental isolates by using cluster methodologies. Furthermore, we show how cluster performance is evaluated using index scores (Silhouette, Calinski-Harabasz, Davies-Bouldin) to ensure the outcome of colony classification. As automation becomes more prominent in microbiology, tools as ColFeatures will assist identification of bacterial colonies on agar plates, bypassing human bias and complementing sequencing or mass spectrometry information that often comes attached with a considerable price tag.

## Introduction

After more than a century of use, the agar plate is still at the core of microbiology (1,2). From counting colony-forming units to surveying microbial diversity, the agar plate remains a cheap and versatile tool. Since each colony on an agar surface originates from a single viable cell, the plate provides both a quantitative measure of the cells added and serves as a viability filter since a colony will only appear if the cell was capable of division.

In this age of big data, analysis of agar plates still provides ample room for improvement. When investigating environmental microorganisms, growth rate, size, morphology, and pigmentation are distinctive features providing crucial information when choosing which colonies to isolate. In addition, these morphological features are helpful to describe phenotypic switching or mutations (3,4). Because the features are often subjective and difficult to document, the processes are often augmented by employing 16S rRNA gene sequencing or mass spectrometry (1,5). This provides more information when selecting colonies or analyzing microbial composition, but comes with a considerable price tag or requires expensive and specialized equipment. Losing chemical diversity from secondary metabolites is an additional caveat of relying on 16S rRNA. Several studies have shown that biosynthetic gene clusters can vary considerably even within strains deemed close to identical by 16S (6–8). Because of this, we hypothesized that digitalization and automation could enable us to use morphological features in a fast and reliable manner to identify and characterize environmental microorganisms.

Counting the number of colonies is relatively simple to automate, and several algorithms have been published (9–15) and some are included in the software for commercial equipment (16). Image segmentation methods are diverse and many different approaches have been implemented. For example, CellProfiler (17,18) is an open-source tool that allows the automation of biological-image analysis, and its modular design allows the development of pipelines to analyze different types of data. More specialized software exists, designed explicitly for colony counting. OpenCFU (10) offered the possibility to identify any type of circular objects from digital pictures using a three-step process. The IMJ Cai Fiji macro (14) relies on edge detection combined with Gaussian smoothing and filling holes to identify bacteria colonies. Zhu and coworkers (12) relied on near-infrared light pictures and developed an automatic four-step method to count colonies. These approaches are reliable for identifying bacteria colonies, but only OpenCFU goes beyond colony identification and retrieves information such as area, perimeter and RGB values.

To provide the microbiology community with an easy-to-use tool to extract morphological information from a single picture, we introduce ColFeatures. Our tool uses a high-resolution image of an isolation plate as input. The computational result is the identified colonies and information on a large set of 28 morphological features for each colony. Furthermore, given that each colony has its ID, colonies can be tracked over time. The user-friendly interface allows the user to fine-tune parameters and provides basic data processing. To demonstrate the potential of our tool, we show how it can be used to cluster bacteria with similar phenotypes and how morphological features can be tracked over time. In summary, ColFeatures represents a new tool in microbiology to automate the study of morphological features in a non-biased way.

## Materials and methods

### Bacterial cultures

Soil collected at Mols Bjerge National Park, Denmark, was used in this study. 1 g of soil was added to 10 ml 0.9% (w/v) saline solution and vigorously vortexed for 10 minutes to create a soil slurry. The soil slurry was allowed to settle, and a ten-fold dilution series was prepared from the supernatant. 50-200 µL of dilutions (10^−4^ or 10^−5^) were plated on 0.1 strength Nutrient Agar (0.1 NA: 0.1 g/L D(+)-glucose, 1.5 g/L peptone, 0.6 g/L NaCl, 0.3 g.l^-1^ yeast extract, 15 g/L agar, pH 7.5) supplemented with nystatin (50 µg/mL). The plates were incubated for 30 days at 22 °C. In the case of our example experiment, soil bacteria strains (both different and similar morphological features – 5 of each) were grown in a liquid medium to an OD_600_ of ∼0.4. The cultures were pooled, a ten-fold dilution series was made in 0.9% (w/v) saline solution. Dilutions were plated on 10% NBA agar plates and incubated for about five days at room temperature.

### Image collection

Images of the agar plates were taken every five days throughout the incubation period using a ProtoCOL 3HD (Synbiosis, Cambridge, UK). The exposure time was adjusted to 190 ms. Plates were accurately positioned to the exact location in the scope to enable temporal tracking of bacterial colonies.

### Clustering

We used k-means clustering, which partitions *n* data points into *k* groups or clusters with equal variance by minimizing the within-cluster sum-of-squares distances. The algorithm takes a value for k given by the user. We used three different metrics to determine the best number of clusters: Silhouette Coefficient score (19), Calisnki-Harabasz score (20) and Davies-Bouldin score (21). The Silhouette Coefficient has a range between -1 and +1; the higher the score, the denser and more separated the clusters are. In the case of the Calisnki-Harabasz Index, dense and well-separated clusters compute a higher score. The third index, Davies-Bouldin Index, compares the distance between the clusters - the closer to zero, the better the partition.

### Cluster strength

We calculated cluster strength based on the spatial distribution of each data point along the 2-component PCA. We first calculated the cluster centroid as the average of all the (x, y) data points that belong to a cluster, only considering clusters with more than two data points. Then, we calculated the distance of each point to the centroid of the cluster and use it to calculate what we defined as cluster strength, *s*.

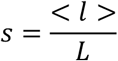

where < *l* > is the average distance between the centroid of the cluster and the data points that belong to that cluster, and *L* is the average distance from a data point to their cluster center across all clusters. Small cluster strength values mean more condense data points inside the cluster, while the larger the cluster strength values, the less packed the data points inside the cluster.

### Software development and distribution

ColFeatures is available from https://github.com/danielaags/ColFeatures. It was implemented in MATLAB^®^ 2019b in Windows 10. Image Processing and Signal Processing Toolbox are required and used as described below. For users with more MATLAB experience, we have included all scripts.

### Image processing workflow

We designed a sequence of steps that allow the computer to pin-point colonies on plates and extract morphological information from them. The workflow is presented in Fig 1.

**Fig 1.**
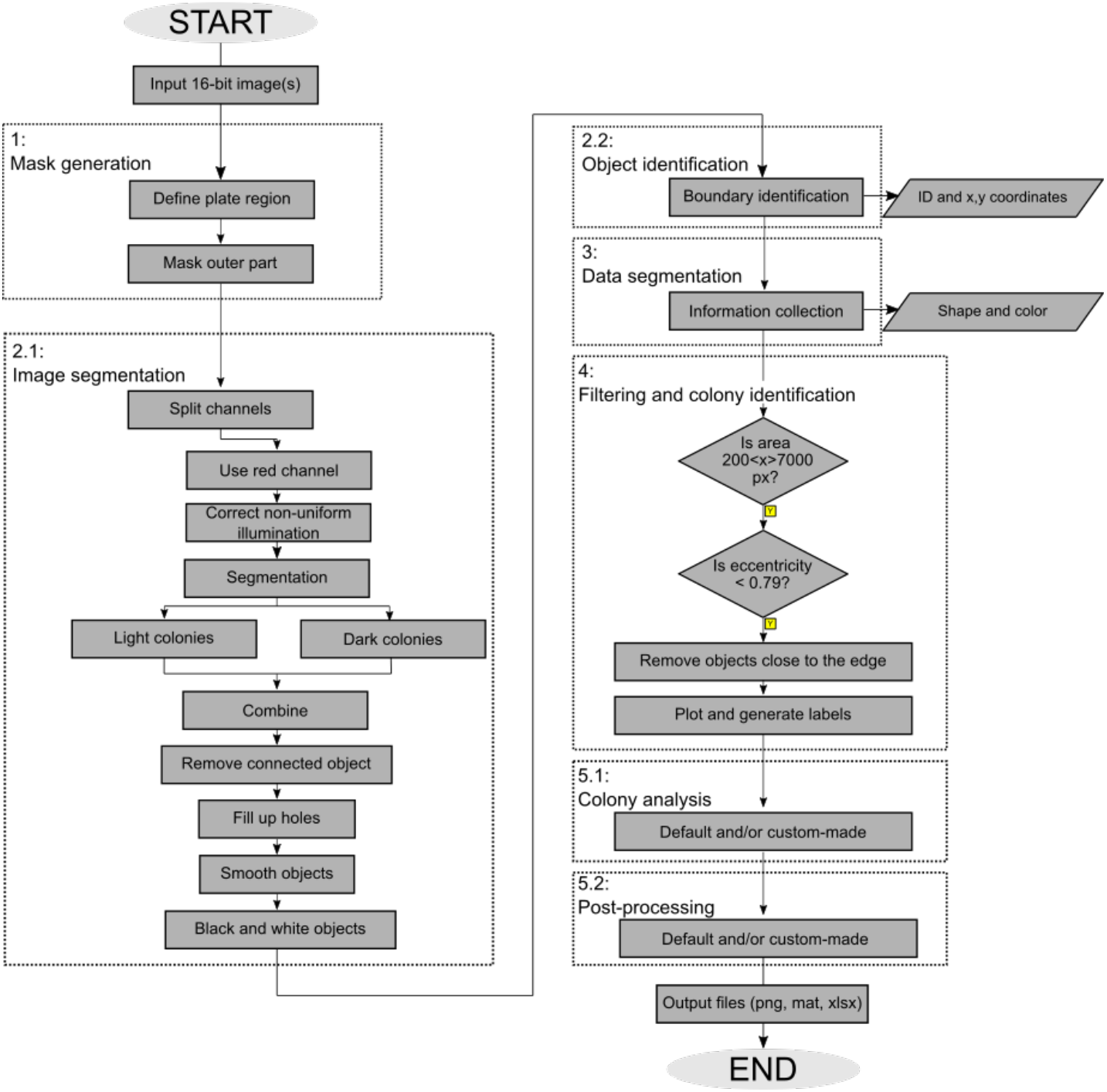
Flowchart representing the main steps implemented. A mask is generated to hide the outer part of the plate (step 1), then segmentation (step 2.1) is done in two steps using only the red channel; after this, object identification (step 2.2) and data segmentation (step 3) is performed. Next, objects are filtered based on the area and eccentricity values (step 4); after this, the remaining objects are identified as colonies, each with a defined label. In the final step, colony analysis and data post-processing (step 5) are carried out either automatically in our GUI or a custom-made analysis depending on the user’s questions.

Our 5-step approach takes a 16-bit image and generates a binary image for labeling and data extraction. We first remove the region outside the agar plate to reduce contrast noise (Step 1). Next, we correct non-uniform illumination and do segmentation in two steps (2.1). We use the segmentation information to define each object (2.2) and extract morphological and pixel intensities (3). Next, we filter the data and identify colonies (4) and do colony analysis (5.1) and post-processing (5.2). Our method can generate an image file with labels for all the colonies identified and create a data file with 36 columns containing morphological and pixel information for each colony.

#### 1. Mask generation

The first step in our algorithm is to convert our 16-bit image into an 8-bit image, with a standard RGB range of 0-255. After this, we remove the outer part of the agar plate and the edge of the plate leaving only the region of interest for our analysis (Fig 2B).

**Fig 2.**
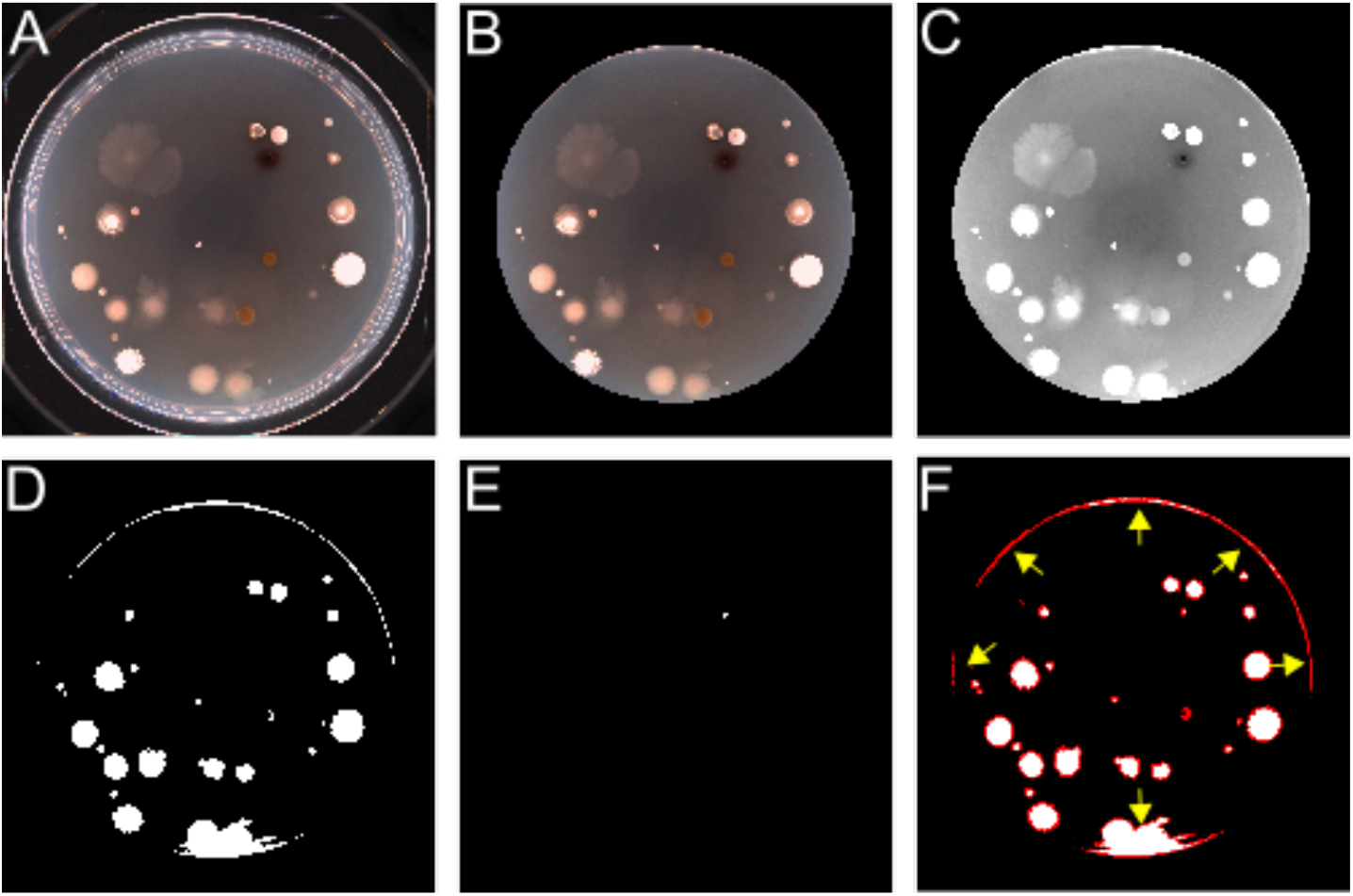
Representative snapshots of the main steps for colony detection. A) Original image. B) Mask generation that removes the outer part of the plate. C) Correction of non-uniform illumination in the red channel. D) Colonies brighter than the background are segmented. E) Colonies darker than the background are segmented. F) After both segmentation outputs are merged, object identification is made, each white independent area is considered an object. A final filtering step is required because some of the identified objects are not bacterial colonies or bacterial colonies with too much noise (yellow arrows).

#### 2. Image segmentation and object identification

Next, we realized that the three RGB channels were not equally informative for colony recognition. After testing a set of images, we decided to use only the red channel for image segmentation. First, we correct for non-uniform illumination to improve the segmentation process (Fig 2C). We noticed two types of colonies in our samples: colonies brighter than the background and colonies darker than the background. Therefore, two steps are required to identify all the colonies on the agar plate, one segmenting bright colonies (Fig 2D) and the other segmenting dark colonies (Fig 2E). After this - to improve the segmentation process - we remove all the connected objects, fill up holes and smooth the objects. The resulting binary black-white (BW) images are used for boundary identification. Here, each colony is identified as an object with a defined ID and a defined position in space. By merging our two-step segmentation process’s boundary information, we got the spatial map of all the colonies (and other artifacts) present on the agar plate (Fig 2F).

#### 3. Data segmentation

We used a built-in MATLAB® function to extract morphological properties and pixel information for each colony. We decided to extract 28 properties that we believe are informative.

### Morphology

We automatically retrieved the following information: centroid, area, diameter, perimeter, circularity, eccentricity and convex hull. Centroid information will give the (x, y) coordinates at which the colony center is located. Area, diameter and perimeter values are given in pixels. Circularity and eccentricity are two values related to the shape of the objects; circularity specifies the roundness of the objects, while eccentricity values could determine if the object is circular (= 0) or a line (=1). Finally, convex hull retrieves the smallest convex polygon inside the boundary. We used the convex hull values combined with the centroid to approximate the topological landscape along the perimeter to calculate the number and height of peaks of each colony. We considered three types of margin: lobate, undulate and entire (S1 Fig). If a bacterial colony has a lobate perimeter, it will have many height peaks and sharp valleys; a bacterial colony with an undulate margin will have fewer and smaller peaks than a colony with a lobate margin. Finally, a bacterial colony with an entire margin will have just a few peaks, and this will be the smallest compared with bacterial colonies with undulate and lobate margin.

### Pixel light intensity information

We extracted the information in the RGB space and Lab space for each colony; we calculated the mean value of each and the standard deviation of the whole area inside the boundaries. We also calculated the mean and standard deviation of the transversal section across each colony.

#### 4. Colony filtering and identification

Since some of the objects segmented in step 2 are background noise, a two-step filtering is applied to keep only objects that are considered bacterial colonies. First, we filtered based on size and shape. We decided to identify only colonies within an area range of 200 – 7000 pixels and an eccentricity value above 0.79. These metrics will retrieve almost all of the colonies seen on the plates, even those with irregular (non-circular) shapes. In a second step, we removed objects that are too close to the rim of the agar plate since these objects were either false positives (noise) or objects where only partial colonies were detected. All the extracted data is filtered and stored in a single data type. We finally plotted the identified colonies on the original image and generated labels for tracking purposes.

#### 5. GUI, post-processing and output files

We developed a user-friendly graphical interface (GUI) that incorporated the described algorithm and allowed for simple colony analysis. The user only needs to load the image to be analyzed and give five different inputs. The inputs defined as the date of experiment and plate number are used to generate labels, the output field is used to name the output files. We also included an option to specify the size of the plate (0 = 90 mm plates, 1 = 150 mm plates) and an extra option to modify the threshold value for segmentation (bright and dark colonies). In this way, the user can increase the identification accuracy of the algorithm. Once input parameters are provided, the user only needs to press ‘Run identification’. The software will generate three different files: a .png file with the identified colonies and the generated labels, a .mat file, and an excel file with all the extracted data together with the generated labels (Fig 3, Identification panels).

**Fig 3.**
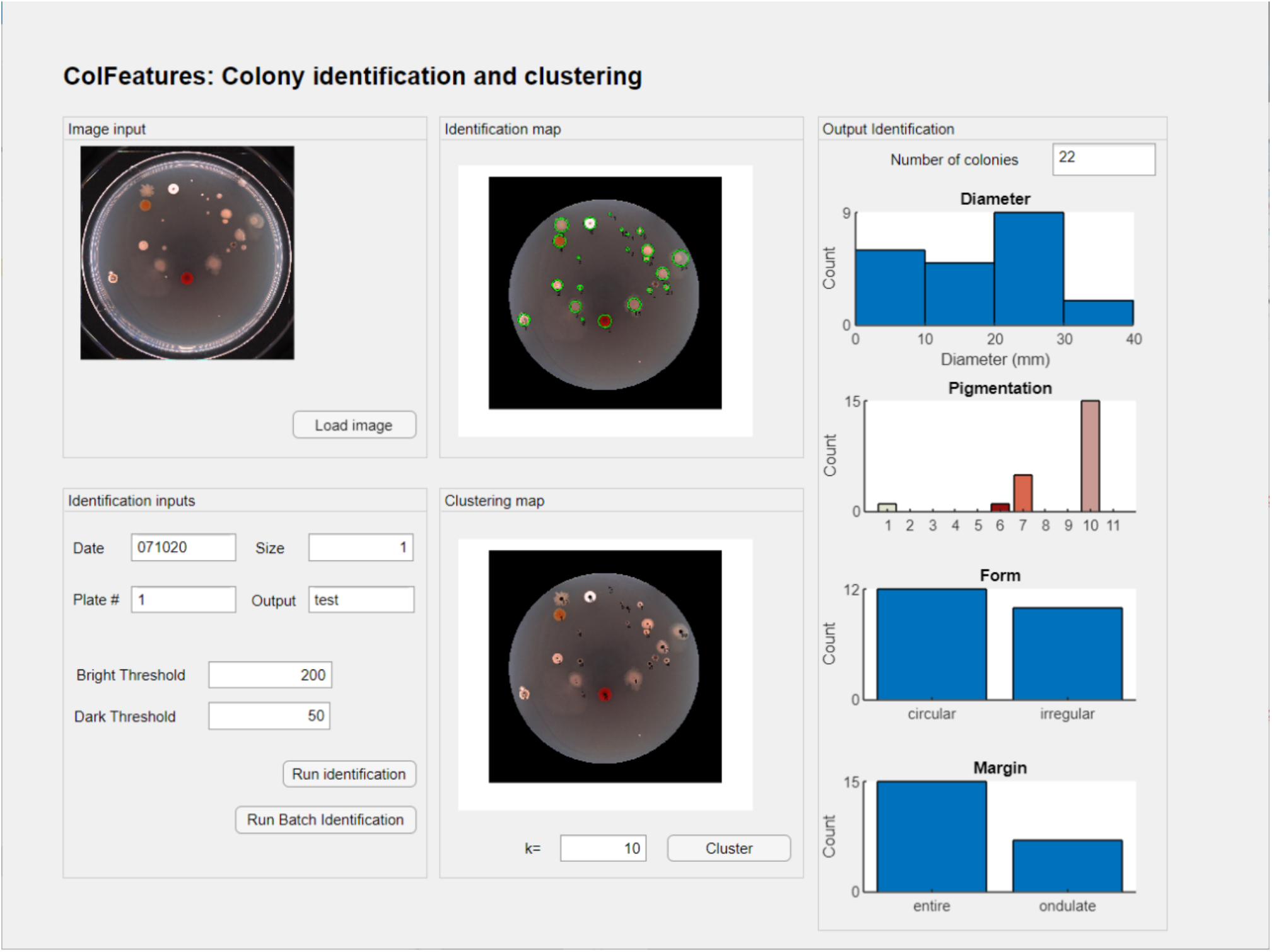
GUI from the described algorithm. Snapshot of the output after clicking ‘Run identification’ followed by ‘Clustering’. After giving the required inputs, the GUI will show the colonies identified with the respective label and display four different plots with standard metrics used in microbiology. The ‘Clustering’ option requires a user-defined number of clusters.

We also included some standard metrics used by microbiologists (diameter, pigmentation, form and margin (4)). First, we displayed the number of colonies that belong to each category. For pigmentation, we used the color scheme reported in Sousa *et al*. 2015: white (1), black (2), grey (3), brown (4), yellow (5), red (6), orange (7), green (8), blue (9), pink (10) and violet (11). Then, by using the RGB mean values and a nearest neighbors’ algorithm, we classified the colonies seen on a plate under this eleven define groups. For form classification, we used the circularity feature and a stringent threshold. Only when circularity == 1 will a colony be defined as circular; else, it will be classified as irregular. Finally, for margin classification, we used the peaks’ height feature with a cutoff threshold of 55 (based on our data). If the height of peaks along the perimeter is >= 55, a colony is classified as undulate, else as entire. All the metrics mentioned above are also included in the .mat and Excel files for further use.

Since we envisioned using morphological features to lead the way to automatic colony classification, we included an additional option to do informed clustering using all the morphological features extracted by our algorithm. The user, based on the experiment, will define the number of clusters to use by typing the value for *k* (Fig 3, Clustering panel). The GUI will then label each colony with its corresponding cluster (Fig 3, Clustering panel). This step will generate a new .png file and a new .mat and excel file with the cluster information. We also included an Identification Batch option. Once the user is satisfied with the threshold values, the batch option will use these values to analyze all the images stored in a defined folder.

## Results

### Colony classification in defined clusters

As a proof of concept, we evaluated our GUI by analyzing artificial examples of isolation plates. We decided to have two different examples: i) one that produced five similar bacterial colonies, ii) and one with five different bacterial colonies. In the example where five similar bacteria were used, a non-trained eye will have trouble seeing the different morphologies (S2A Fig). After using our identification and clustering option (Fig 4A and 4B), we noticed that most of the colonies were identified. Next, to better visualize how these colonies were classified based on the morphological features, we used PCA to reduce the dimensionality of our data and plot using only two dimensions. Given the high similarity between the colonies, we observed that the five expected clusters are not well defined and have considerable overlap (Fig 4C). When looking at the cluster strength (s) values, the cluster with the lowest s’ values (cluster five) has more evenly distributed data points than the one with the lowest s’ (cluster two), which also seems to have more outliers. Moreover, the PCA shows that given the bacterial morphological features, it is more likely to have two rather than five clusters since the data points partition in one cluster at the left-upper region of the PCA and a second cluster in the right-bottom region.

**Fig 4.**
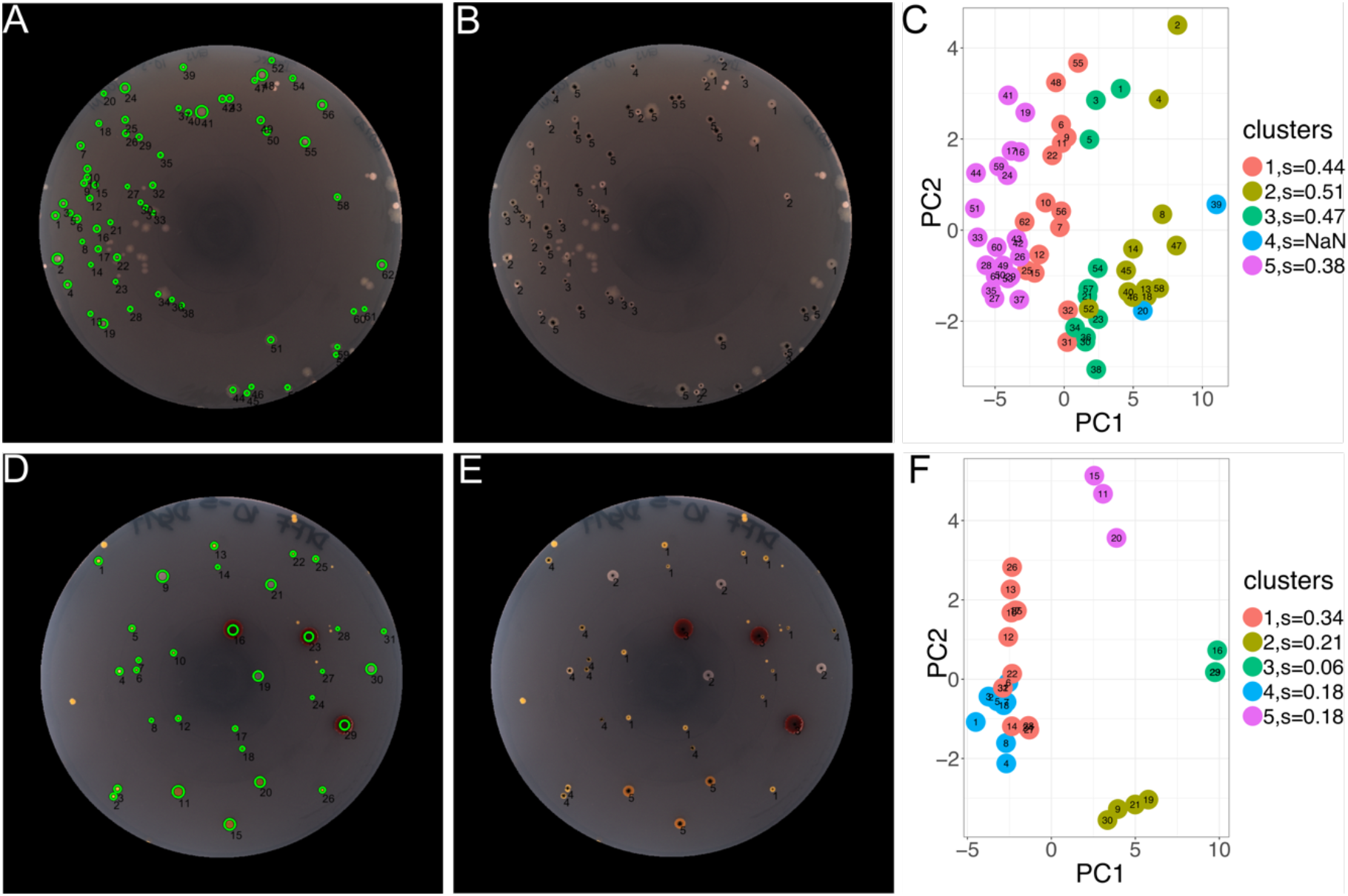
Colony identification and clustering of artificial examples of isolation plates. A) Representative snapshot of an experiment where five very similar bacterial colonies were plated. The labeled bacterial colonies after identification and data extraction and B) clustering classification are shown. C) Two-component PCA for data visualization, the corresponding clusters are plotted in a specific colour. D) Representative snapshot of an experiment where five different bacterial colonies were plated. The labeled bacteria colonies after identification and data extraction and E) clustering classification are shown. F) Two-component PCA for data visualization, the corresponding clusters are plotted in a specific color.

When looking at the data from the example where five different colonies were plated (S2B Fig), the colonies were more distinctive - even to an untrained eye - and the algorithm had fewer problems identifying and clustering the colonies (Fig 4D and 4E). The 2-component PCA analysis shows clustering with better-separated clusters (Fig. 4F) than the previous example (Fig 4C) and the clusters are more compact. However, if we consider clusters zero and three, it seems that the bacterial colonies might belong to the same cluster. This example shows how our tool can help with the identification and clustering of bacterial colonies from agar plates.

### Tracking bacterial colonies over time

In our following example, we decided to test our tool in batch. We used 16 images from agar plates with soil bacteria from two different isolation rounds. We were interested in investigating the change of morphological features over time. We picked two data points: 15 days and 20 days and decided to use a subset of eight plates per time point. We used k-means clustering (22) to partition our data based on 28 different variables. Since performance evaluation is not trivial, we decided to determine the best number of clusters to partition our data using three different metrics that work on data where the ground truth labels are unknown. Since each metric calculates k-means performance differently, we reasoned that combining the three metrics to determine the *k* value or the number of clusters, the better it will be supported. The three metrics used were the following: Silhouette Coefficient score (19), Calisnki-Harabasz score (20) and Davies-Bouldin score (21) (See Material and Methods). We evaluated the cluster performance with k=2 to k=212 (total number of data points) and observed the inflection point and best values for the three different score/indexes when k=5 (Fig5A – red dotted line). Overall, when comparing the data from 15 days against the data from 20 days, we see that at 20 days, the clusters have a better partition as seen by the three different metrics used (Silhouette Coefficient score: 15 days = 0.27 and 20 days = 0.28; Calisnki-Harabasz Index score 15 days = 86 and 20 days = 76 and Davies-Bouldin Index score 15 days = 1.5 and 20 days = 1.3) (Figs 5B).

**Fig 5.**
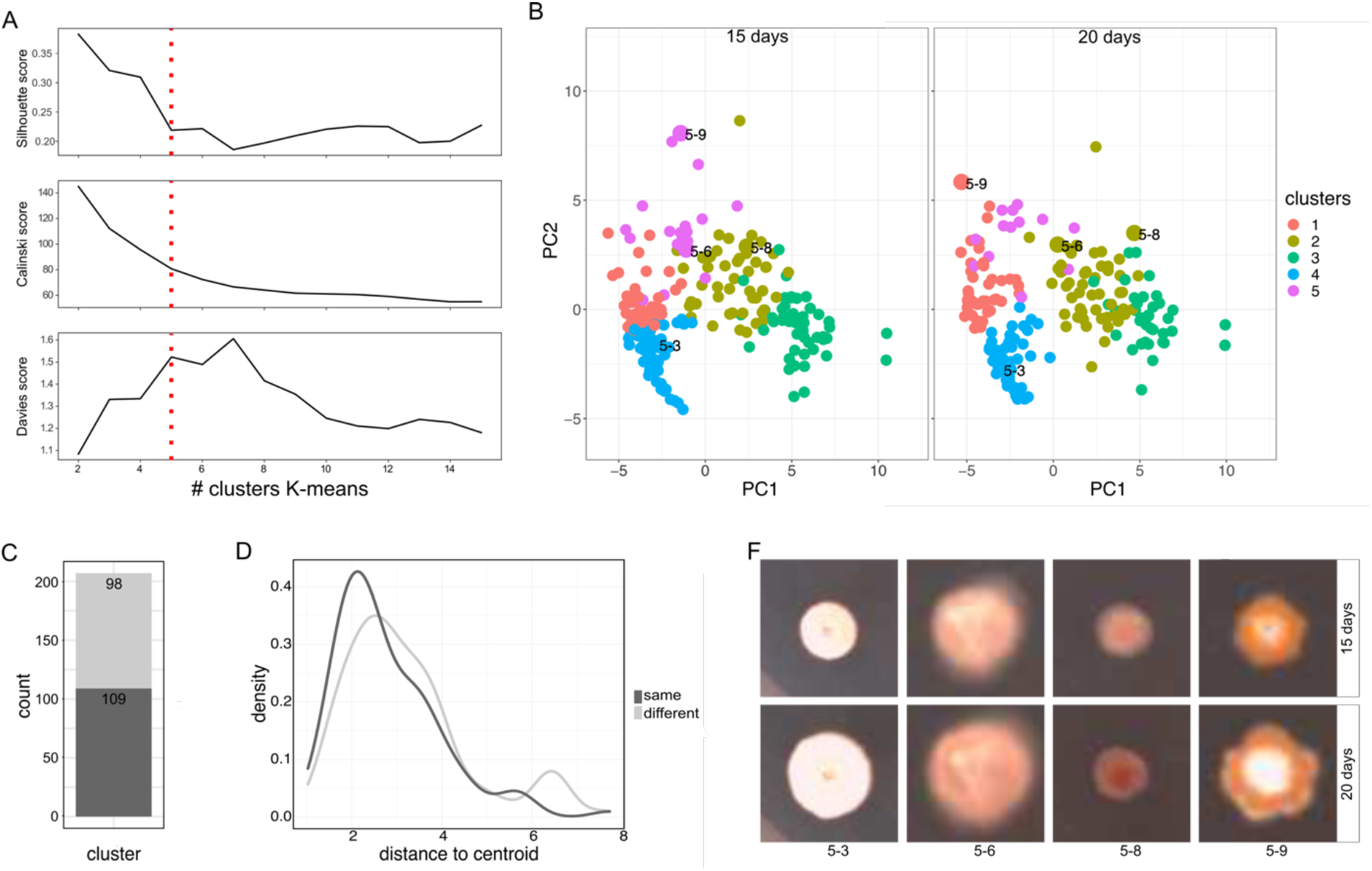
Data partition of bacterial colonies in five clusters at two different time points. A) Subset of 212 bacterial colonies, k values in a range 0-15 and the different values obtained for each metric. We see the inflexion point in k = 5. B) The 2-component PCA and classification of bacterial colonies in five clusters at 15 days (left-panel) of incubation and 20 days (right-panel) of incubation. Four bacterial colonies are set as examples. C) Bar plot showing the proportion of bacterial colonies that remain in the same cluster (dark gray) and those that move to a different cluster (light gray) from data at 15 days and 20 days. D) Distribution of the distance between the cluster centroid and the data point from colonies that remain in the same cluster (distance _mean_ = 2.8) and colonies that move to a different cluster (distance _mean_ = 3.1). Higher distance values to the centroid of the cluster are expected for colonies that move to a different cluster after 20 days of incubation. E) Example of bacterial colonies that remain in the same cluster (5-3 and 5-8) and an example of bacterial colonies that move to a different cluster (5-6 and 5-9). For a complete picture of one of the agar plates used, see S3 Fig.

Interestingly, the proportion of bacterial colonies that move to a different cluster between the two-time points was almost half, supporting the idea that morphological features are changing even after a long incubation period (Fig 5B). It seems that such property can be easily tracked by determining the distance between a data point and the centroid of the cluster that they belong to (Fig5D and 5E); the larger the distance, the more likely for a data point to move to a different cluster.

## Discussion

We have introduced the software ColFeatures which identifies and retrieves morphological information from bacterial colonies on agar plates. Colony identification is carried out using a two-step segmentation, ensuring that colonies brighter and darker than the background are identified. Furthermore, since the appearance of bacteria colonies is a widely used tool by microbiologists (2), our algorithm offers the possibility to automatically extract information that describes the appearance of bacterial colonies on agar plates, improving our ability to document colony morphology automatically in a consistent manner.

Our software has two different implementations that will depend on the user’s needs. In any case, the main outputs will be: 1) an image with all the colonies identified, each with its label, and 2) a data frame with all the morphological information. Using the GUI implementation, the user can explore the software and have some examples of how the automatically extracted information can be used and interpreted. At the same time, the user can explore the possibility of clustering bacterial colonies based on morphological properties. In one of our examples, ColFeatures GUI could easily identify and cluster bacterial colonies. We observed that overall automatic classification is less time-consuming and more consistent when comparing automatic and manual clustering. If the user has a large image dataset, we recommend using the batch mode to speed up the analysis.

ColFeatures shows several advantages over manual description. The most important is how fast and easy it is to extract morphological features from colony plates, which would be very time-consuming even for an experienced microbiologist. Another significant advantage of our software is that it will allow consistency in the data and unbiased comparison between different samples, researchers, and experiments. However, we believe there is still room for improvement. As an example, our software works best for light agar plates. We also explored using dark agar plates (S4 Fig), but found colony identification quite challenging. Pixel quality and illumination are essential factors to take into consideration while collecting the data. It is important to use pictures with high pixel quality; the better the quality, the more likely to have meaningful morphological information from each image. Illumination is paramount when comparing different experiments. One needs to use the same settings such as exposure time; otherwise, comparison between different samples is impossible. When tracking changes along the time, during image acquisition, it is quite essential to place the plates in the same position, in our case we decided to draw a mark on one edge of the plate and align it to a marked place on the plate holder. It is also highly recommended to work with samples with few fused bacteria since the software cannot deal with such data.

We believe that this method represents a valuable tool for microbiologist, as it is easy to use and capable of automatically extracting important morphological information either alone or in combination with sequencing or mass-spectrometry data.

