## Supporting Information for "ColFeatures: Automated data extraction and classification of bacterial colonies"

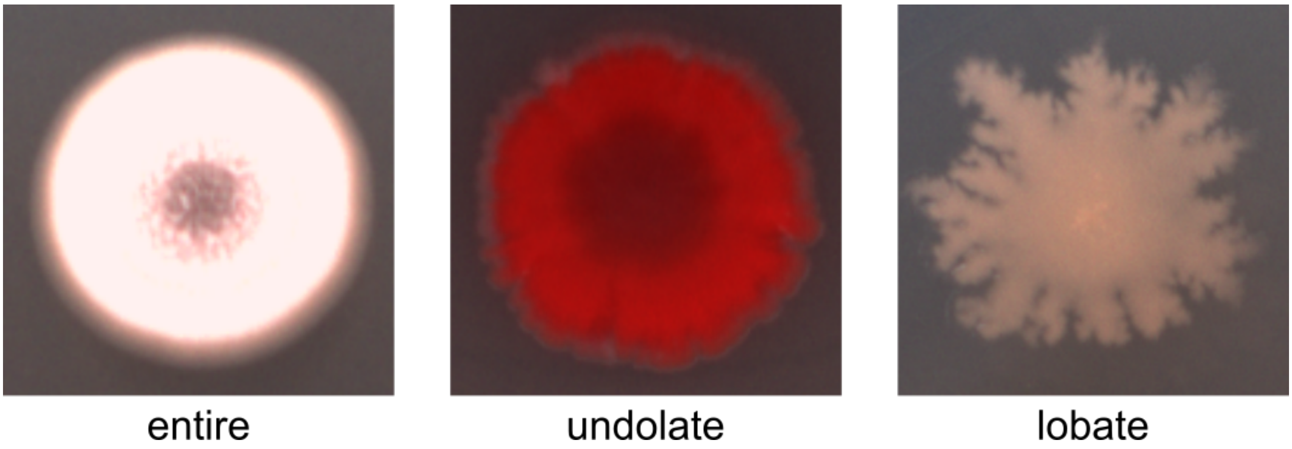


**S1 Fig. Colony margin classification.** Three different types of margin can be define based on the peaks/valleys around the edge of a bacterial colony.

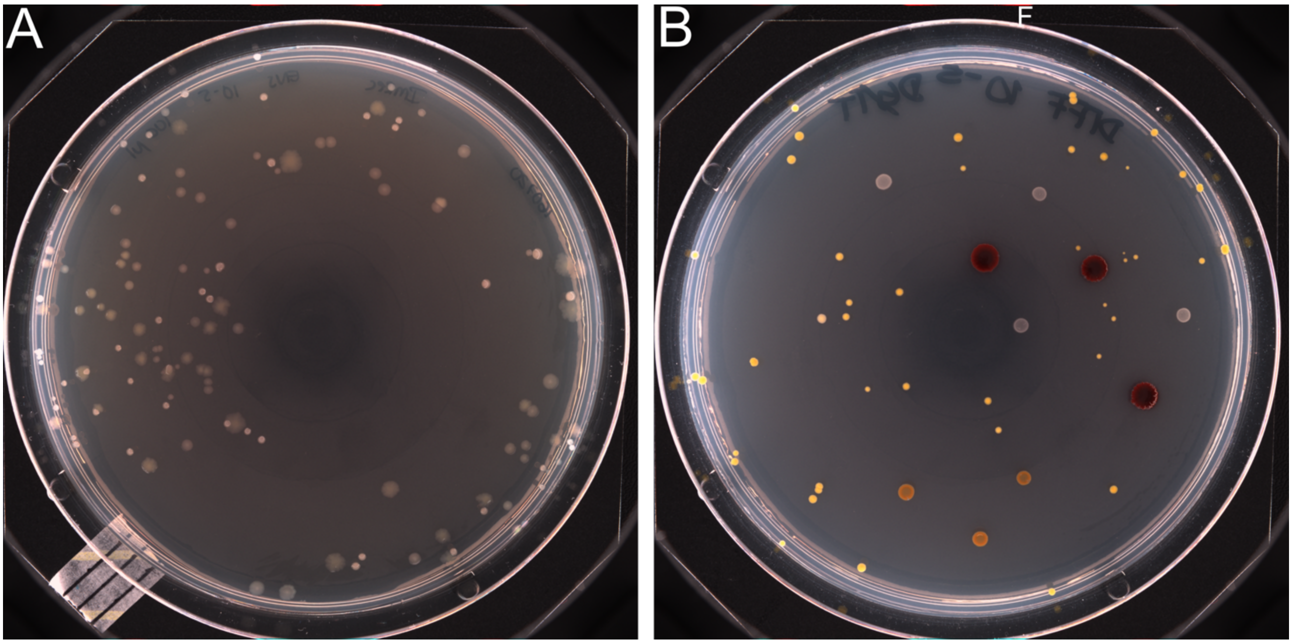


**S2 Fig. Artificial examples of isolation plates.** A) Snapshot of an agar plate from an experiment where five very similar bacterial colonies were plated. B) Snapshot of an agar plate from of an experiment where five very similar bacterial colonies were plated.

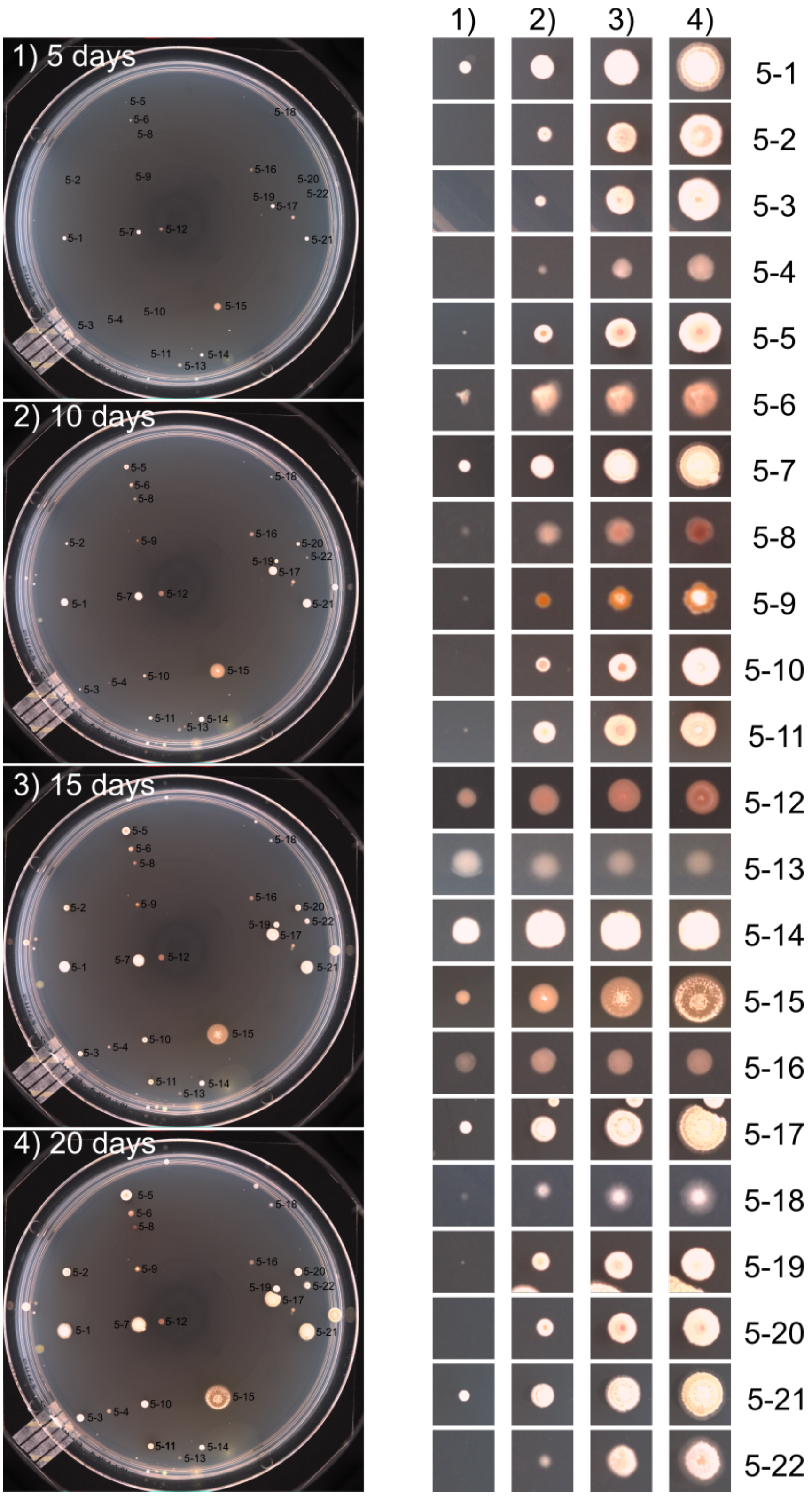


**S3 Fig. Example of one of the sixteen agar plates tracked. Pictures were taken every 5 days for a total of 20 days.** The bacterial colonies are labeled with their ID on the four data points. One can notice that some of the colonies take longer to be visible than others **(**left panels). Thumbnails of the colonies tracked are shown (right panels).

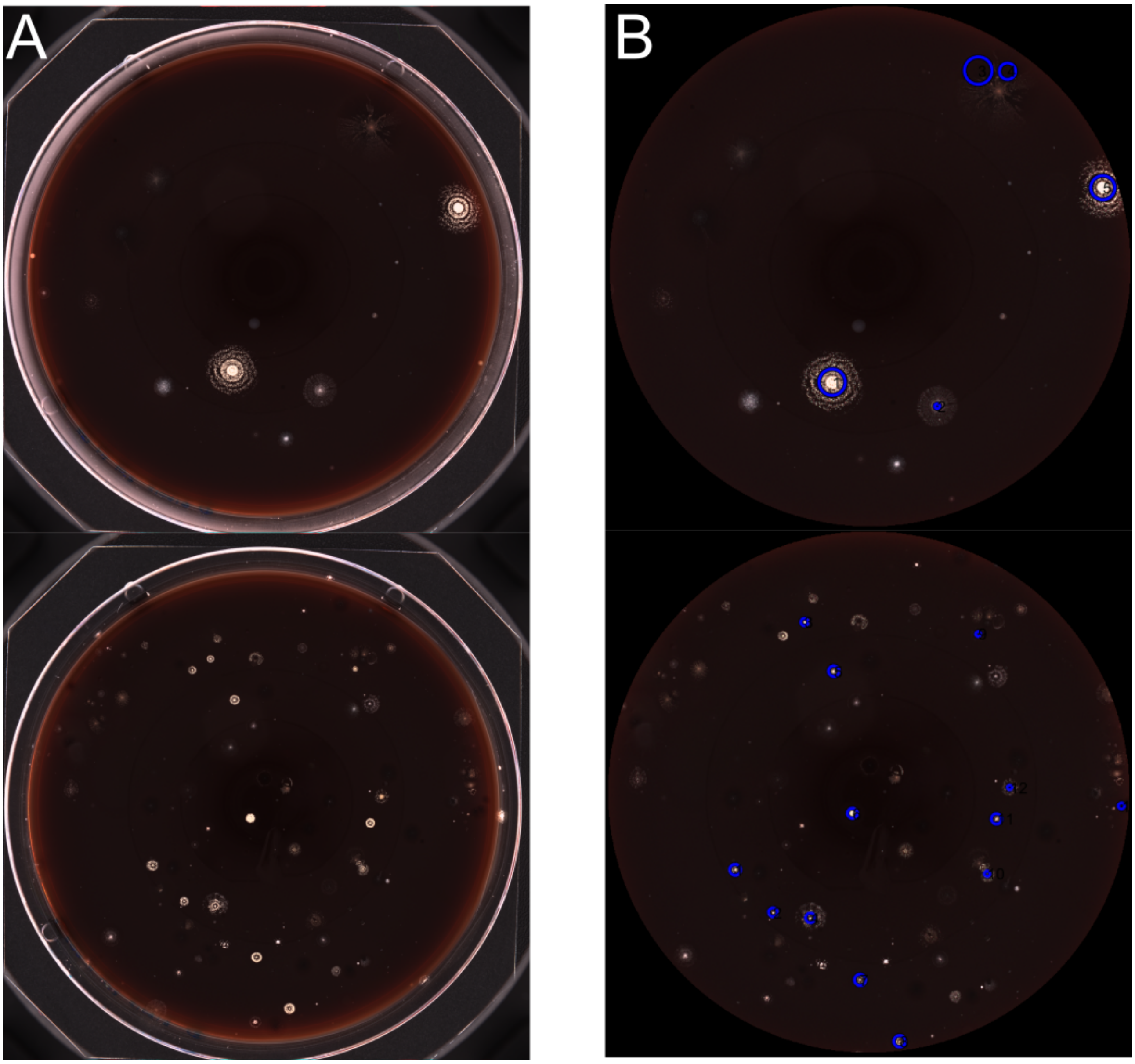


**S4 Fig. Examples of agar plates with a dark background.** A) Representative pictures of HAV agar plates after incubation with soil bacteria for 30 days at 22 C. B) Output of ColFeatures from plates in panel A, the algorithm as troubles to identify each colony on the plate.
